# *In Vivo* Screening for a Promising Antiparasitic Agent Against *Neobenedenia melleni* in *Epinephelus fuscoguttatus*♀ × *E. lanceolatus*♂ and Identification of Its Potential Target

**DOI:** 10.64898/2026.08.24.746648

**Authors:** Longkun Gao, Guangshuo Wang, Juntao Xu, Yanru Guo, Wei Luo, Ying Yan, Guanhai Li, Qin Yu, Mingzhu Liu, Erlong Wang, Pengfei Li, Tianqiang Liu

## Abstract

Monogenean ectoparasites, particularly *Neobenedenia* species, cause severe economic losses in mariculture. Here, ectoparasites isolated from cultured hybrid groupers (*Epinephelus fuscoguttatus*♀ × *E. lanceolatus*♂) were confirmed as *Neobenedenia melleni* based on ITS1 phylogeny. *In vivo* screening of six structurally diverse compounds identified compound D (CAS No. 206111-37-7), a 5,6-dihydropyridine derivative, as the most effective antiparasitic agent, achieving 76.54% efficacy at 0.5 mg/L in a 90 min bath treatment. Dose-response assays demonstrated that 0.7 mg/L compound D achieved 95.23% antiparasitic efficacy without causing evident tissue damage or cytotoxicity to GF-1 cells. Ultrastructural observation by scanning electron microscopy revealed marked tegumental alterations, including deep fissures and extensive surface folding, in treated parasites. Molecular docking against ten candidate proteins identified β-tubulin as the most favorable docking target, with a binding energy of −6.53 kcal/mol and three hydrogen-bond interactions, suggesting that β-tubulin may be involved in the antiparasitic activity of compound D. Overall, these findings highlight compound D as a promising lead candidate for short-bath therapy against *N. melleni* and suggest that cytoskeletal disruption through β-tubulin interaction represents a plausible mechanism of action.

## 1. Introduction

Marine aquaculture is now a major source of aquatic food worldwide, supporting food security and coastal economies [1]. However, the expansion of intensive farming has also increased the impact of parasitic diseases, which can cause substantial production losses each year [2,3]. Among these parasites, *Neobenedenia* spp. (Monogenea: Capsalidae) are important ectoparasites of marine fishes because of their broad host range and considerable pathogenicity [4–6]. They have been reported from a wide range of hosts, including commercially important reef-associated and pelagic fishes such as groupers, snappers, butterflyfishes, and mullets [7–9]. In particular, *Neobenedenia melleni* has been recorded from more than 100 host species belonging to approximately 30 families, indicating an exceptionally broad host range for a monogenean parasite [10,11].

*Neobenedenia* spp. primarily inhabit the skin, fins, and eyes of their host fish, where they attach to the epithelial surfaces using a specialized haptor equipped with hooks [12]. Heavy infections can cause excessive mucus production, epithelial erosion, and ulceration, which compromise host condition and may increase the risk of secondary infections [13,14]. In severe cases, infected fish may show reduced feeding and growth, and mortality can occur, resulting in considerable economic losses [7,15]. These pathogenic effects, together with the high stocking densities used in intensive aquaculture, favor parasite transmission and rapid population increase, making effective control of *Neobenedenia* infections an important challenge for marine fish production.

The biological characteristics of *Neobenedenia* spp. also make their control difficult. They have a direct life cycle without an intermediate host, and their eggs hatch into free-swimming oncomiracidia that actively seek and attach to suitable hosts [16]. Temperature and other environmental conditions strongly influence parasite development and reproduction [17]. Under favorable conditions, *Neobenedenia* spp. can complete their life cycle rapidly and produce large numbers of eggs, allowing infections to become established and persist in aquaculture systems [18,19]. Their eggs are also less susceptible to some control measures than actively attached parasites and may remain in the culture environment, providing a source of reinfection [20]. Thus, controlling established parasites alone may not be sufficient to prevent recurrent infections.

Chemotherapy is widely used to control *Neobenedenia* infections, and praziquantel (PZQ) is one of the most extensively studied anthelmintic agents against monogeneans [21,22]. PZQ can markedly reduce *Neobenedenia* burdens under suitable treatment conditions, but its efficacy varies with factors such as drug concentration, exposure duration, treatment regimen, and parasite developmental stage. Several studies have reported strong activity against adult *Neobenedenia*, whereas its effect on parasite eggs is generally limited [23,24]. Freshwater or hyposalinity treatments and chemical disinfectants can also reduce parasite burdens, but their application may be constrained by host tolerance, treatment conditions, environmental concerns, or limited activity against less susceptible developmental stages. These limitations have prompted interest in alternative antiparasitic agents that can complement existing control measures and improve the management of *Neobenedenia* infections.

Screening compounds with known or potential bioactivity offers a practical approach to identifying new agents against *Neobenedenia*. Phenotypic screening is particularly useful for an initial evaluation because parasite responses such as changes in motility, detachment, viability, and morphology can be assessed directly [25]. Previous studies have shown that this approach can identify compounds with activity against *Neobenedenia* spp., including plant extracts and purified plant-derived compounds with substantial effects on adult parasites [26,27]. Such findings suggest that compounds from different chemical classes may provide useful candidates for further evaluation. However, compared with parasitic diseases of medical and veterinary importance, relatively few studies have systematically evaluated structurally diverse compounds against marine monogeneans [28]. Moreover, promising compounds identified in initial screening still require concentration-dependent evaluation and testing in host fish to determine their antiparasitic efficacy and safety.

In this study, we identified the *Neobenedenia* species infecting cultured groupers using morphological examination combined with ITS1 sequencing and phylogenetic analysis. We then evaluated a panel of compounds with different chemical structures for their antiparasitic activity against the parasite *in vivo*. The most active compound was subsequently evaluated over a range of concentrations to determine its antiparasitic efficacy and tolerance in the host fish. Its effects on the parasite surface were also examined by scanning electron microscopy to provide morphological evidence of drug-induced damage. Finally, molecular docking was used to compare the interactions of the selected compound with several parasite-associated proteins, with particular attention to proteins previously implicated in parasite survival or antiparasitic drug action. Through this approach, we aimed to identify a promising compound for the control of *Neobenedenia* infection and provide preliminary evidence for its possible molecular target.

## 2. Materials and methods

### 2.1 Parasites collection

Parasites were collected from the body surface of *Epinephelus fuscoguttatus*♀ × *E. lanceolatus*♂ cultured at a fish farm in Zhangzhou, Fujian, China. The fish were anesthetized with 0.02% MS-222 (tricaine methanesulfonate, Aladdin Scientific, Shanghai, China) according to the manufacturer’s instructions. Parasites were carefully removed from host skin using forceps and a scalpel, preserved in 70% ethanol, and examined under an Olympus BX53 microscope.

### 2.2 DNA extraction, amplification, and sequencing

Approximately 10 parasites were collected from the body surface of the groupers for molecular identification. Genomic DNA was extracted using the TIANamp Marine Animals DNA Kit from TIANGEN (TIANGEN Biotech, Beijing, China). The internal transcribed spacer 1 (ITS1) region of the ribosomal RNA gene was amplified by polymerase chain reaction (PCR) using the primers F (5′-GTCGTAACAAGGTTTCCGTAGG-3′) and R (5′-GCTGCACTCTTCATCGACGCRCG-3′). PCR amplification was performed with an initial denaturation at 94 °C for 5 min, followed by 35 cycles of denaturation at 94 °C for 30 s, annealing at 55 °C for 30 s, and extension at 72 °C for 1 min. A final extension was performed at 72 °C for 10 min. The purified PCR products were subjected to bidirectional Sanger sequencing by Sangon Biotech (Shanghai, China). The resulting sequences were edited and assembled, followed by BLAST searches against the NCBI GenBank database for sequence similarity analysis.

### 2.3 Phylogenetic analysis

Except for the ITS1 sequences obtained in the present study, all sequences used for the phylogenetic analysis were retrieved from the NCBI GenBank database. The sequences of *N. melleni* (HQ684821.1, HQ684822.1, HQ684823.1, HQ684826.1, HQ684830.1, HQ684831.1, HQ684832.1, HQ684835.1, HQ684837.1, HQ684838.1, HQ684842.1, AY551323.1), *N. girellae* (AY551326.1, JF934745.1), *Dactylogyrus ctenopharyngodonis* (KX369213.1), and *Dactylogyrus dulkeiti* (KX369217.1) were included in the phylogenetic analysis as reference sequences. The best-fit nucleotide substitution model was determined using ModelFinder based on the Bayesian information criterion (BIC), and the T92 + G model was selected for subsequent phylogenetic analysis in MEGA 12.

### 2.4 Fish preparation and *N. melleni* challenge

Commercial groupers (*E. fuscoguttatus*♀ × *E. lanceolatus*♂, 109 ± 28 g) were subjected to freshwater immersion for 30 s to remove ectoparasites before the experiment. The fish were subsequently acclimated in holding tanks for 2 weeks. During acclimation and subsequent experimental procedures, seawater temperature was maintained at 28 ± 0.5 °C, dissolved oxygen was maintained above 6 mg/L, pH was 7.6 ± 0.3, and salinity was maintained at 27-29 ‰. The seawater was recirculated through an ultraviolet sterilization system and a protein skimmer. Fish were fed commercial pelleted feed twice daily at a daily feeding rate equivalent to 2% of body weight. Experimental infection with *N. melleni* was performed using a protocol modified from Hirazawa et al [7]. Briefly, uninfected recipient fish were cohabitated with parasitized donor fish for a period of 7 days. After removal of the donor fish, the recipients were maintained for an additional two weeks before subsequent experiments.

### 2.5 *In vivo* antiparasitic efficacy assay

Groupers infected with *N. melleni* were individually transferred into separate 20 L experimental tanks containing filtered seawater pre-adjusted to the corresponding drug concentrations. Fish were immersed for 90 min, with continuous aeration to maintain dissolved oxygen. The number of *N. melleni* attached to the body surface of each fish was counted and recorded immediately before and after the immersion treatment. Antiparasitic efficacy was calculated using the following formula: Antiparasitic efficacy (%) = (N_0_ − N_1_) / N_0_ × 100%, where N_0_ represents the number of parasites attached to the fish surface before treatment, and N_1_ represents the number of parasites remaining on the fish surface after treatment. All compounds used for the screening experiments were purchased from Aladdin (Shanghai, China). Detailed information on the tested compounds is provided in Table 1.

**Table 1.** Information on the compounds tested for antiparasitic efficacy against *N. melleni*.

| Compound | CAS No. |
| --- | --- |
| triclabendazole | 68786-66-3 |
| matrine | 519-02-8 |
| mycophenolic acid | 24280-93-1 |
| 1-(tert-Butyl) 3-ethyl 4-hydroxy-5,6-dihydropyridine-1,3(2H)-dicarboxylate<br>(compound D) | 206111-37-7 |
| tropolone tosylate | 38768-08-0 |
| artemisinin | 63968-64-9 |

Based on the preliminary screening results, compound D was further evaluated at six concentrations (0.1, 0.2, 0.3, 0.5, 0.6, and 0.7 mg/L), with three fish per group (n = 3) and a control. Infected groupers were immersed in filtered seawater containing the corresponding concentrations of compound D for 90 min under continuous aeration. The number of remaining *N. melleni* was recorded after treatment, and the antiparasitic efficacy was calculated as described above.

### 2.6 Toxicity assessment of compound D

The acute immersion toxicity of compound D to groupers and its cytotoxicity toward Grouper Fin-1 (GF-1) cells (PTA-859, purchased from ATCC Bioresource Center) were evaluated. Healthy groupers were randomly divided into a control group and six treatment groups exposed to compound D at concentrations of 0.1, 0.3, 0.5, 0.7, 1.0, and 2.0 mg/L, with three fish in each group. Fish were immersed for 90 min in 20 L tanks under continuous aeration, and water parameters were consistent with Section 2.4. Fish survival was recorded after exposure.

The cytotoxicity of compound D toward GF-1 cells was evaluated using the Cell Counting Kit-8 (CCK-8) assay. GF-1 cells were seeded in 96-well plates and cultured in L-15 medium supplemented with 10% fetal bovine serum (FBS) at 28 °C. After cell attachment, the cells were exposed to serially diluted concentrations of compound D and incubated for 48 h. Blank and control groups were included, with three replicate wells for each concentration. Following drug exposure, CCK-8 reagent was added to each well, and the plates were incubated in the dark for 3 h at 37 °C. Absorbance was subsequently measured at 450 nm using a microplate reader. Cell viability was calculated using the following formula: Cell viability (%) = (A_t_ − A_b_) / (A_c_ − A_b_) × 100%. where A_t_ represents the absorbance of the treatment group, A_b_ represents the absorbance of the blank group, and A_c_ represents the absorbance of the control group.

### 2.7 Histopathological examination

Following the 90 min immersion assay at 0.7 mg/L compound D (maximum tolerated concentration) and 0.2% DMSO solvent control, groupers were euthanized with an overdose of 0.02% MS-222. Tissue samples including heart, liver, spleen, and kidney were dissected immediately. Tissues were fixed in 4% paraformaldehyde for at least 24 h at room temperature. After fixation, the samples were subjected to gradient ethanol (30%, 50%, 70%, 85%, and 95%) dehydration, xylene clearing, and paraffin embedding, followed by sectioning at a thickness of 5 μm. The sections were stained with hematoxylin and eosin (H&E), mounted on glass slides, and examined under a light microscope.

### 2.8 Scanning electron microscopic (SEM) analysis

Untreated and compound D-treated *N. melleni* were prepared for SEM analysis. The treated parasites were exposed to compound D at 0.7 mg/L for 90 min, while untreated parasites served as controls. Following treatment, the parasites were collected and fixed in 2.5% glutaraldehyde at 4 °C overnight. The samples were washed three times with phosphate-buffered saline (PBS) and dehydrated sequentially in 30%, 50%, 70%, 85%, and 95% ethanol, each for 15-20 min, followed by two changes of 100% ethanol for 20 min each. The samples were subsequently treated three times with 100% tert-butanol for 30 min each and subjected to critical point drying. After gold sputter coating, the ultrastructural morphology of untreated and compound D treated *N. melleni* was examined using a Hitachi SU8600 scanning electron microscope.

### 2.9 Molecular docking analysis

Potential target proteins were selected for molecular docking analysis. Protein annotation and sequence matching information from a previous proteomic study of *Neobenedenia* sp. were used to identify the corresponding proteins and sequence information [29]. For cathepsin B (GW923086.1), glutathione transferase (GW920036.1), tropomyosin (GW920884.1), and β-tubulin (GW918166.1), the corresponding nucleotide sequences were retrieved from the NCBI Expressed Sequence Tags (EST) database. Open reading frames were predicted using the NCBI ORF Finder, and the deduced amino acid sequences were used for subsequent structural prediction. The amino acid sequences of heat shock protein 70 (CBM69253.1), heat shock protein 90 (CBM69255.1), annexin B1 (ANW06232.1), and cathepsin L-like cysteine protease (ABK62795.1) were obtained from the NCBI Protein database. For actin-1 and calcium-transporting ATPase, the amino acid sequences of *Schistosoma mansoni* (P53470.1) and *Fasciola hepatica* (THD27339.1), respectively, were used. The obtained amino acid sequences were subjected to three-dimensional structure prediction using AlphaFold 3, and the predicted structures were subsequently used for molecular docking analysis. Molecular docking was performed using AutoDock 4 with AutoDockTools (ADT) for receptor and ligand preparation. Briefly, the predicted protein structures were prepared by removing water molecules, adding polar hydrogen atoms, and assigning Gasteiger charges. Compound D was prepared as the ligand, and the docking grid box was defined to encompass the entire protein structure of each target. Docking calculations were performed using the Lamarckian genetic algorithm implemented in AutoDock 4. The docking results were compared to identify the protein showing the most favorable interaction with compound D, and the corresponding protein-compound D complex was visualized using PyMOL 2.4.0.

### 2.10 Statistical analysis

All data were presented as the mean ± standard deviation (SD). Data analysis and graph plotting were performed using GraphPad Prism 8.0.2.

## 3. Results

### 3.1 Parasite identification and phylogenetic analysis

Microscopic examination showed that the collected monogeneans were oval-shaped and exhibited the general morphological characteristics of *Neobenedenia* sp. (Fig. 1A). Genomic DNA was extracted from individual parasites, and the ITS1 region of the ribosomal RNA gene was amplified by PCR. Agarose gel electrophoresis revealed a single clear band of approximately 400 bp (Fig. 1B). The PCR product was subsequently subjected to Sanger sequencing, generating a high-quality ITS1 sequence. The sequence obtained in this study was: TTCCTTACGACTGTACATTGGTTGCATCCTTGCACTAATCCTCTAAAAAATGGGGGCTACGGTCCCACGAGCCCGTCGAGGCTTCACCATTATGGTCCCTAACACCGGTGTACCGCATCGGAAGTGGGGCTGTTCAACTGTTGGATCGCATCTTGCAACTGTGCGTGAAGGCATTTGGTATGTTGGACAGCAATCACATTAGTGAAACCATCATGGGTACGTTGCAAGCCATGAATTGCATTTATTGCAATGAGTAATTTCCTAACTTCTATGTGGCGACGTAAGTCGTCATGTTACAGCGCCCCAGAAATGGGGAGGAAGCTGTAAAACTTTCACTCCATGTGGTGGATCACTCGGCTCGTGCGTCGATGAGAAAGTGCAGC. BLAST searches against the GenBank database identified multiple closely related *Neobenedenia* spp. The ITS1 sequence obtained showed the highest sequence similarity to *Neobenedenia melleni*, with 99.73% nucleotide identity and 91% query coverage. *Neobenedenia girellae* sequence also showed high nucleotide identity (99.70%); however, its query coverage was lower (82%). A Maximum-likelihood phylogenetic tree was constructed based on ITS1 sequences from *Neobenedenia* spp. and outgroup monogeneans (Fig. 2C). The isolate obtained in this study, designated Δ ITS1, clustered firmly within the *N. melleni* clade together with multiple reference *N. melleni* sequences, whereas *N. girellae* formed a distinct branch. Collectively, the BLAST analysis and phylogenetic placement supported the identification of the monogenean parasite recovered from cultured groupers as *Neobenedenia melleni*.

**Fig. 1.**
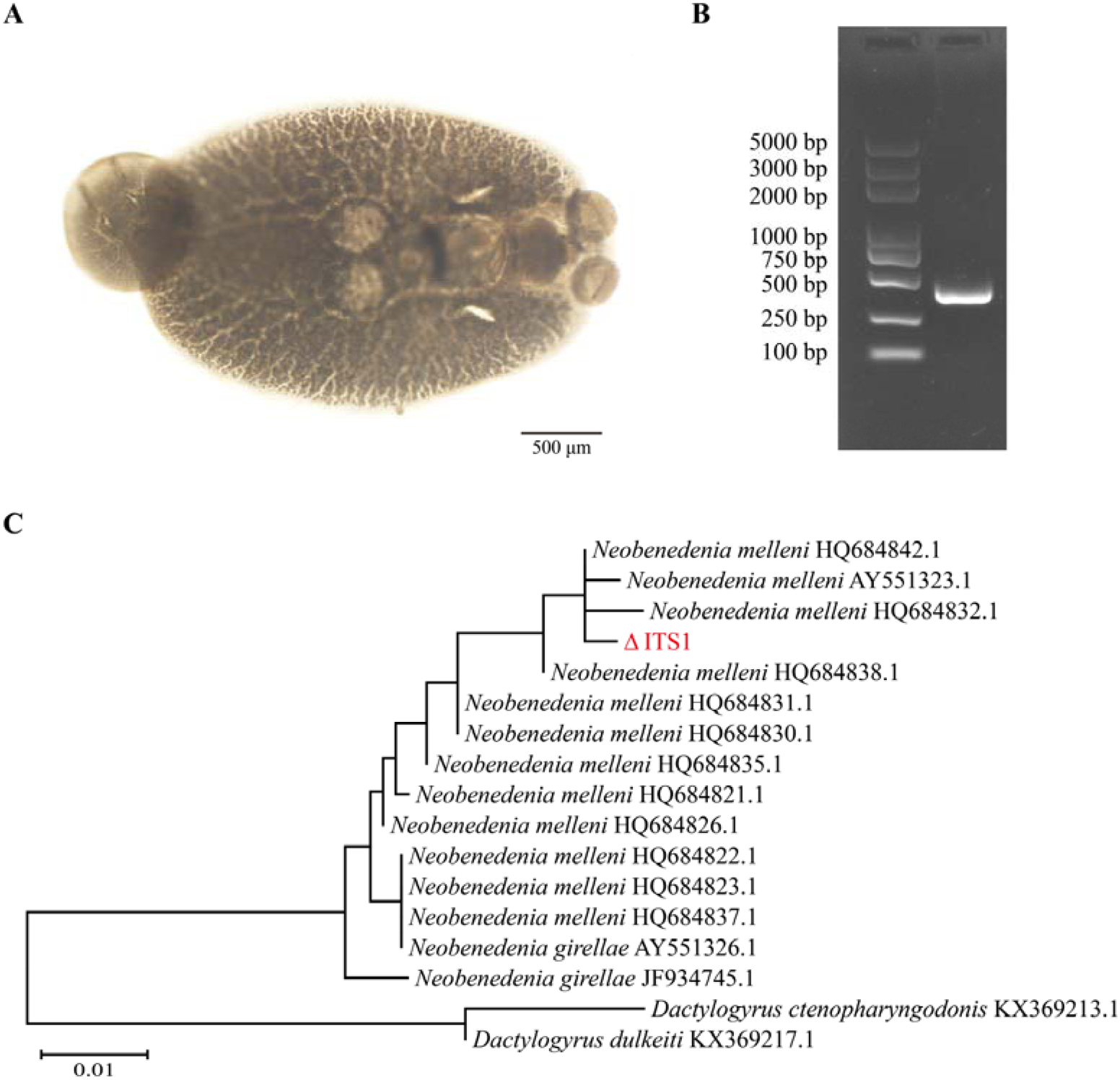
Light micrograph, ITS1 PCR amplification and maximum-likelihood phylogenetic analysis of *N. melleni*. (A) Light micrograph of *N. melleni* collected from an infected grouper. Scale bar = 500 μm. (B) Agarose gel electrophoresis of the ITS1 PCR product. (C) Maximum-likelihood phylogenetic tree based on ITS1 sequences of monogenean parasites. The ITS1 sequence obtained in this study is marked as Δ ITS1. The scale bar represents genetic distance.

**Fig. 2.**
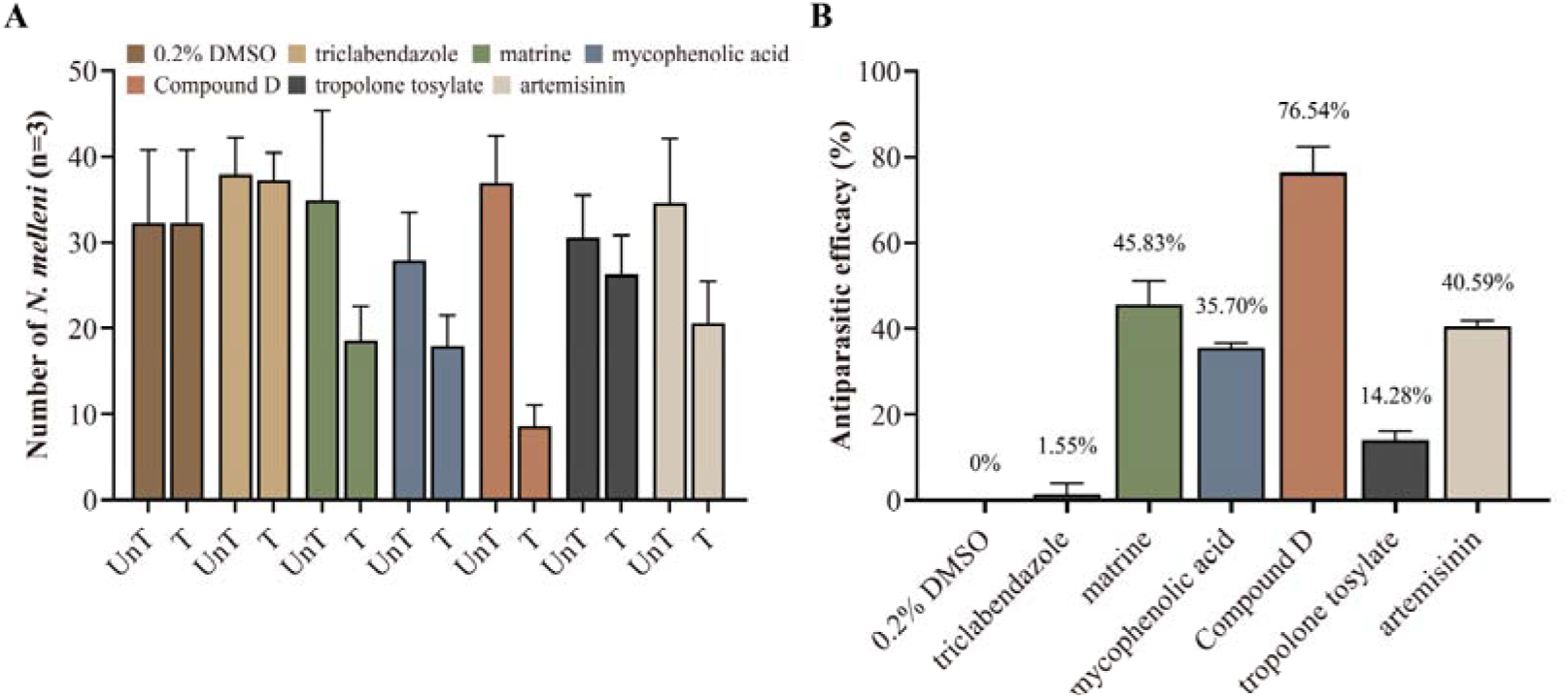
*In vivo* antiparasitic efficacy of candidate compounds (0.5 mg/L) against *N. melleni* in infected groupers. (A) Number of *N. melleni* on fish before treatment (UnT) and after 90 min immersion treatment (T). 0.2% DMSO was used as the solvent control. (B) Calculated antiparasitic efficacy (%) of each compound. Data are expressed as mean ± SD (n = 3).

### 3.2 *In vivo* antiparasitic efficacy of candidate compounds

To identify potential candidates for the control of *N. melleni*, six compounds with diverse chemical scaffolds and reported antiparasitic or related bioactivities were selected and evaluated for their *in vivo* antiparasitic efficacy. All compounds were tested at a uniform concentration of 0.5 mg/L, with 0.2% DMSO serving as the solvent control. Figure 2A shows the parasite burden in infected groupers before treatment (UnT, untreated) and after 90 min of treatment (T, treated). Marked reductions in parasite counts were observed for all compounds except the solvent control and triclabendazole. Subsequently, antiparasitic efficacy was calculated and the results are presented in Figure 2B. At 0.5 mg/L, compound D exhibited the highest antiparasitic efficacy (76.54%), followed by matrine (45.38%) and artemisinin (40.59%). Overall, compound D showed the highest *in vivo* antiparasitic efficacy after 90 min of treatment at 0.5 mg/L and was therefore selected for subsequent evaluation.

### 3.3 Cytotoxicity and acute toxicity of compound D

To further evaluate the safety of compound D, its cytotoxicity against GF-1 cells was determined across a series of concentrations. As shown in Fig. 3A, cell viability remained relatively high at concentrations of 0.1-5.0 mg/L, with 79.05% viability observed at 5.0 mg/L. Cell viability gradually decreased with increasing compound D concentrations and declined to approximately 46% at 50.0 mg/L. These results indicated that compound D exhibited relatively low cytotoxicity at the concentration used for the antiparasitic efficacy assay. The *in vivo* safety of compound D was further evaluated by monitoring the survival of groupers exposed to different concentrations (Fig. 3B). All fish survived at concentrations of 0.1-0.7 mg/L. At 1.0 mg/L, only one of three fish survived, whereas all fish died at 2.0 mg/L. To assess the potential histopathological effects of compound D on host tissues, heart, liver, spleen, and kidney tissues from fish treated with 0.7 mg/L compound D for 90 min were subjected to H&E staining (Fig. 3C). Histological examination revealed no obvious pathological alterations, with no signs of necrosis, inflammatory infiltration, hemorrhage or tissue degeneration, when compared with the 0.2% DMSO solvent control group. Collectively, these results indicated that compound D was well tolerated by groupers at concentrations up to 0.7 mg/L under the tested acute immersion conditions.

**Fig. 3.**
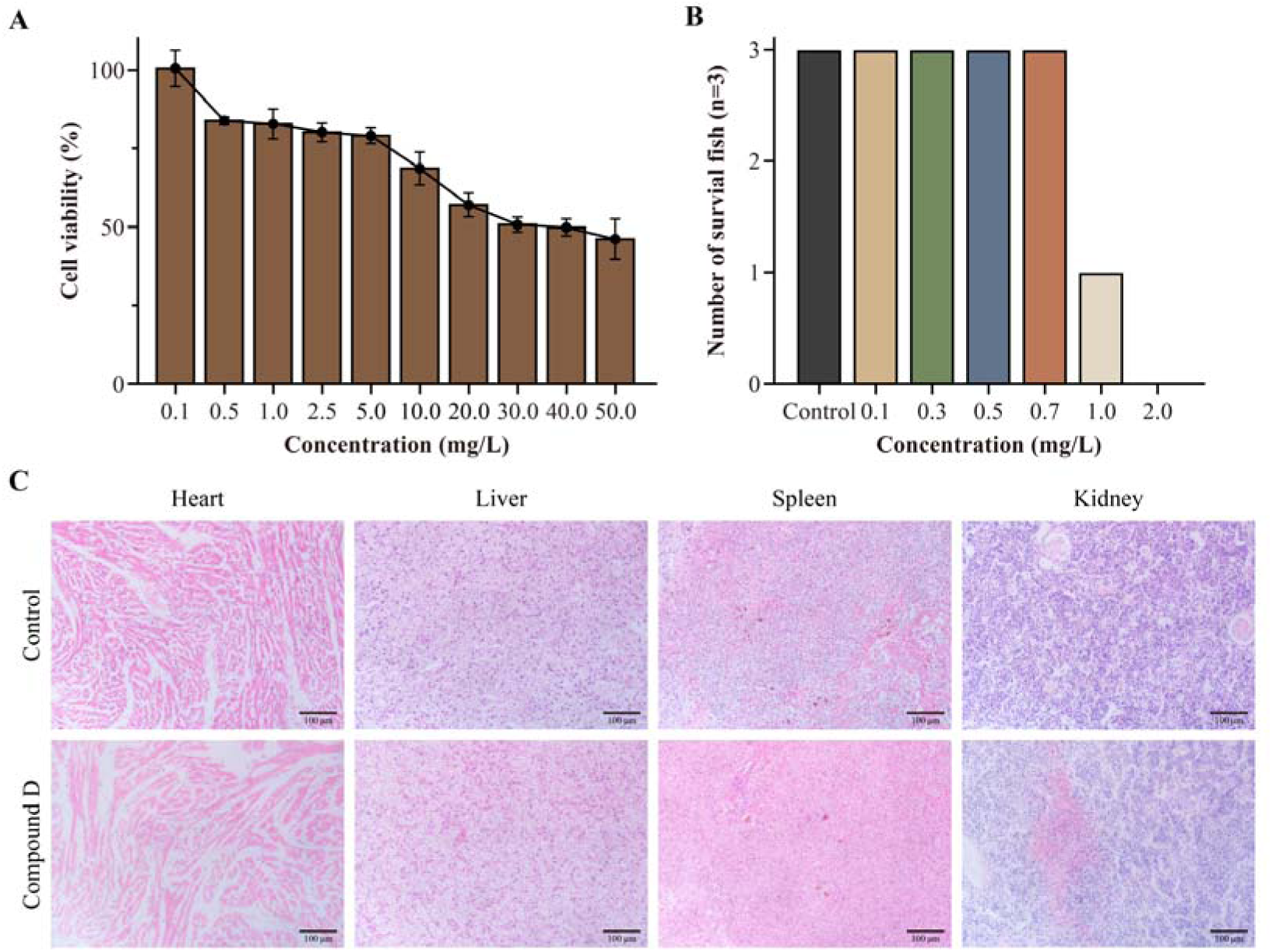
Cytotoxicity, acute immersion toxicity and histological evaluation of compound D. (A) Viability of GF-1 cells following exposure to gradient concentrations of compound D. Data are expressed as mean ± SD (n = 3). (B) Survival number of groupers after immersion treatment. 0.2% DMSO served as solvent control. (C) H&E-stained histological sections of heart, liver, spleen and kidney from fish in the solvent control group and 0.7 mg/L compound D treated group. Scale bars = 100 μm.

### 3.4 Concentration-dependent antiparasitic efficacy and ultrastructural damage of *N. melleni* induced by compound D

To further characterize the antiparasitic efficacy of compound D, its concentration-dependent efficacy was evaluated, and SEM was performed to examine ultrastructural alterations in *N. melleni*. As shown in Fig. 4A, compound D produced a clear concentration-dependent increase in antiparasitic efficacy over the tested concentration range. The antiparasitic efficacy reached 76.54% at 0.5 mg/L, and further increased to 95.23% at 0.7 mg/L. SEM images revealed obvious morphological differences between control and compound D exposed parasites. Parasites in the 0.2% DMSO solvent control group exhibited an intact surface structure with a smooth and regular surface (Fig. 4B, C). In contrast, parasites treated with 0.7 mg/L compound D showed severe surface damage, characterized by extensive surface folding, fissures, and structural disruption across the body surface (Fig. 4D, E). These findings indicated that compound D induced pronounced surface damage in *N. melleni*, which may contribute to its antiparasitic efficacy.

**Fig. 4.**
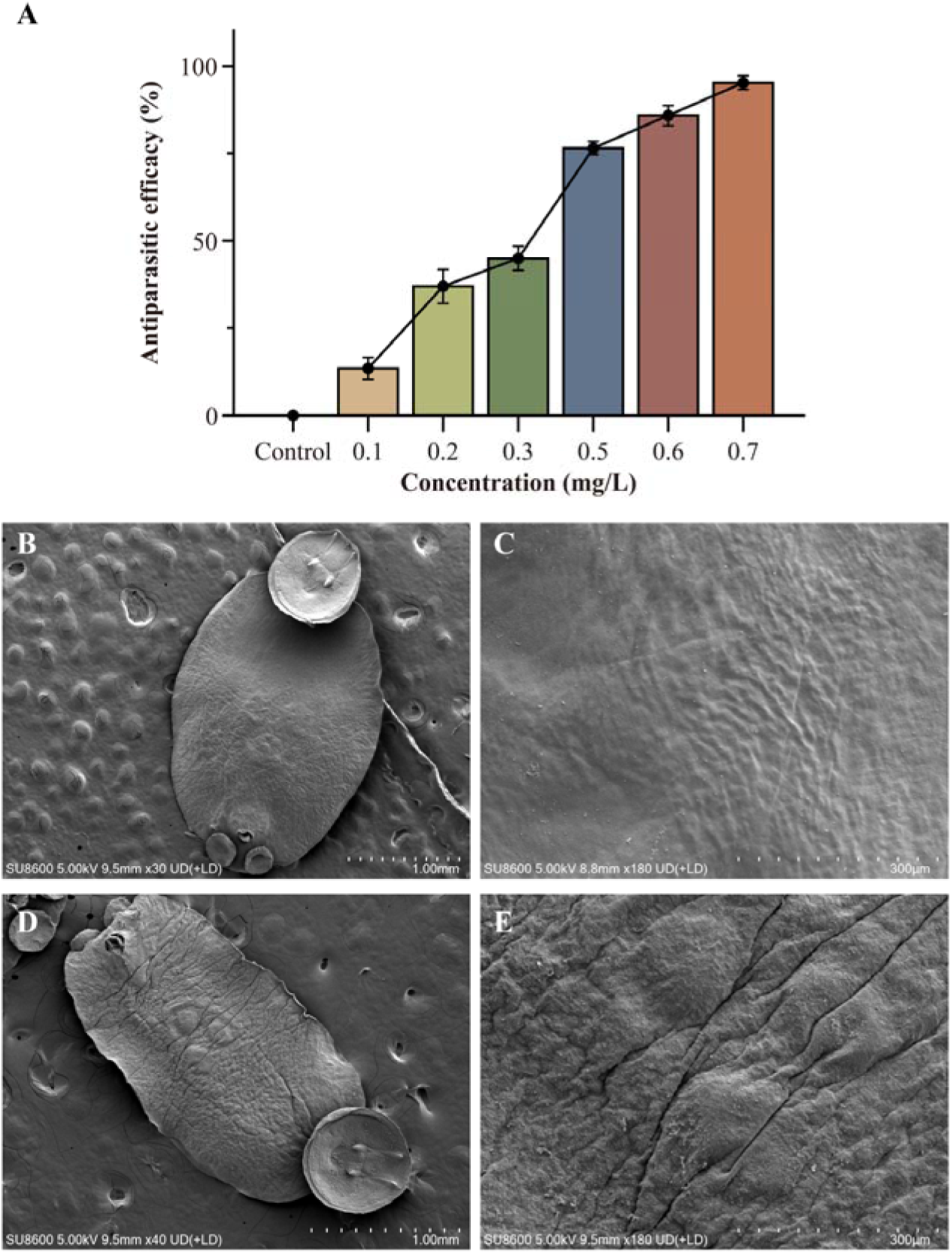
Concentration-dependent antiparasitic efficacy and ultrastructural changes of *N. melleni* following compound D exposure. (A) Antiparasitic efficacy of compound D at different concentrations. 0.2% DMSO served as solvent control. (B-C) SEM micrographs of *N. melleni* from 0.2% DMSO control group. (D-E) SEM micrographs of *N. melleni* treated with 0.7 mg/L compound D. Data are expressed as mean ± SD (n = 3).

### 3.5 Molecular docking analysis of compound D against potential target proteins of *N. melleni*

To explore the potential molecular targets underlying the antiparasitic activity of compound D, molecular docking was performed against ten candidate proteins of *N. melleni*. These potential targets, including cytoskeleton-associated proteins and previously reported drug targets in parasites, were selected based on proteomic studies of monogenean parasites (*Neobenedenia* sp.) [29]. The binding energy, predicted inhibition constant (Ki), ligand efficiency, intermolecular energy, and number of hydrogen bonds for each protein-ligand complex are summarized in Table 2. Among the ten potential targets, β-tubulin exhibited the most favorable overall docking performance. Compound D showed the lowest binding energy toward β-tubulin (−6.53 kcal/mol), corresponding to a predicted Ki of 16.39 μM and a ligand efficiency of −0.34 kcal/mol HA^−1^. The intermolecular energy was also lowest for the β-tubulin with compound D complex (−8.32 kcal/mol). Three hydrogen bonds were predicted between compound D and β-tubulin. In comparison, the other potential proteins showed higher binding energies ranging from −1.44 to −3.67 kcal/mol and substantially higher predicted Ki values ranging from 2.03 to 88.72 mM. Although annexin B1 also formed three hydrogen bonds with compound D, its binding energy (−3.67 kcal/mol) and predicted Ki (2.03 mM) were markedly less favorable than those of β-tubulin. These results indicated that β-tubulin exhibited the strongest predicted binding affinity for compound D among the ten potential proteins. The binding mode of compound D within the β-tubulin binding pocket was further visualized (Fig. 5). Compound D was accommodated within the binding pocket and formed three hydrogen bonds with MET255, ARG297, and LEU299, with distances of 2.2, 1.9, and 1.7 Å, respectively. Collectively, the molecular docking results identified β-tubulin as the potential protein showing the most favorable predicted interaction with compound D, suggesting that β-tubulin may represent a putative molecular target of compound D in *N. melleni*.

**Fig. 5.**
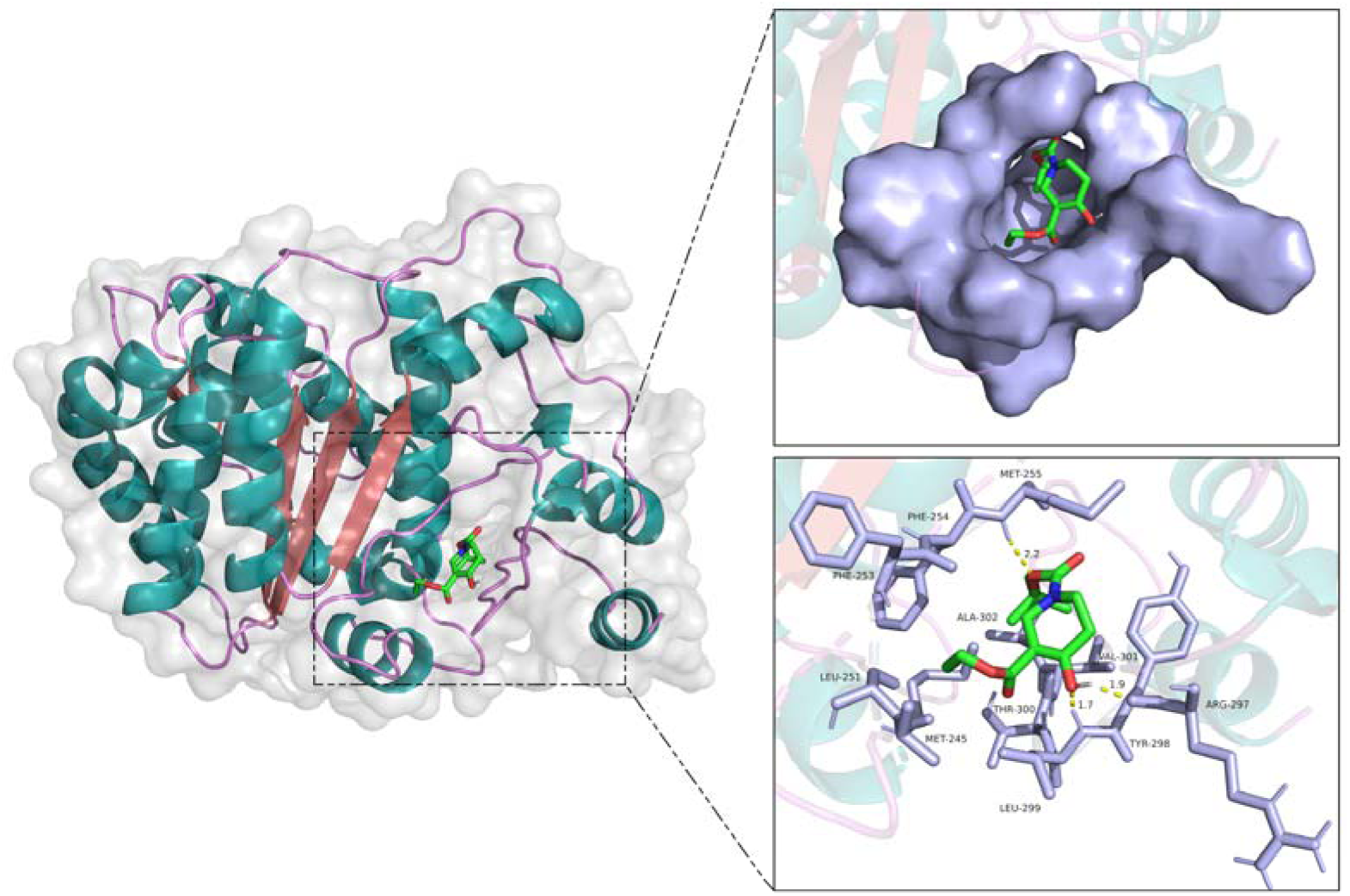
Predicted binding mode of compound D with *N. melleni* β-tubulin from molecular docking. Overall structure of β-tubulin (AlphaFold 3 predicted model) bound to compound D (left panel). Enlarged view of the binding pocket (upper right). Detailed 3D view showing hydrogen-bond interactions and key surrounding amino-acid residues (lower right).

**Table 2.** Binding properties of compound D docked against potential target proteins of *N. melleni*.

| Protein | Binding energy (kcal/mol) | Predicted Ki (μM) | Ligand efficiency (kcal/mol HA<sup>-1</sup>) | Intermolecular energy (kcal/mol) | H-bonds |
| --- | --- | --- | --- | --- | --- |
| cathepsin B | -3.12 | 5,170 | -0.16 | -4.91 | 0 |
| glutathione transferase | -3.61 | 2,260 | -0.19 | -5.40 | 1 |
| tropomyosin | -1.44 | 88,720 | -0.08 | -3.22 | 0 |
| β-tubulin | -6.53 | 16.39 | -0.34 | -8.32 | 3 |
| heat shock protein 70 | -2.34 | 19,150 | -0.12 | -4.13 | 0 |
| heat shock protein 90 | -2.31 | 20,100 | -0.12 | -4.10 | 1 |
| annexin B1 | -3.67 | 2,030 | -0.19 | -5.46 | 3 |
| cathepsin L-like cysteine protease | -2.43 | 16,600 | -0.13 | -4.22 | 0 |
| actin-1 | -2.57 | 13,140 | -0.14 | -4.36 | 2 |
| calcium-transporting ATPase | -1.76 | 51,440 | -0.09 | -3.55 | 0 |

## 4. Discussion

Monogenean ectoparasites represented by *Neobenedenia* spp. constitute a persistent bottleneck for mariculture, causing substantial economic losses via skin injury, secondary bacterial infection and host mortality. Species identification is particularly important for *Neobenedenia* because the taxonomic relationship between *N. melleni* and *N. girellae* has long been debated. Earlier taxonomic revisions considered the two taxa to be conspecific, regarding *N. girellae* as a junior synonym of *N. melleni* based on overlapping morphometrics [10]. However, recent multigene phylogenetic studies have challenged this view and reinstated *N. girellae* as a valid, distinct species [6]. This taxonomic shift, combined with high phenotypic plasticity and host-induced morphological variations [30], renders species identification based solely on morphology highly problematic. In the present study, the collected parasites were first recognized as *Neobenedenia* sp. based on morphology and subsequently delineated using the ITS1 region. The sequence obtained showed the highest similarity to *N. melleni* in BLAST analysis, with higher query coverage than *N. girellae*. More importantly, the isolate clustered with reference *N. melleni* sequences in phylogenetic analysis, with *N. girellae* forming a sister branch. Taken together, these results confirm the identity of our specimens as *N. melleni*, providing a clear taxonomic basis for this study while highlighting the close relationship between these two congeners that allows for meaningful comparison with existing antiparasitic research on both species.

Screening structurally diverse small-molecule libraries against target ectoparasites is a pivotal approach for discovering antiparasitic agents for marine aquaculture [31]. In this study, we evaluated a panel of candidate compounds comprising both benchmark antiparasitic and representative small-molecule with distinct pharmacological mechanisms (matrine, artemisinin, triclabendazole, mycophenolic acid, tropolone tosylate and compound D). The variable success of established antiparasitic agents in short-term immersion therapy underlines how host-parasite interface physiology limits drug efficacy in monogeneans. While matrine reduces *Cryptosporidium parvum* oocyst shedding by 54-63% and restores mucosal barrier integrity in infected mice, as evidenced by decreased plasma D-lactate, reduced bacterial translocation, and lower LDH activity [32], and artemisinin exerts antiparasitic activity via heme-mediated reductive activation of the endoperoxide bridge, generating C-centered radicals that alkylate heme and parasite proteins [33,34], these agents were developed primarily for systemic or oral administration against endoparasites, leaving their efficacy during brief immersion of surface-attached ectoparasites largely unexplored. Likewise, the lack of efficacy of mebendazole against *Microcotyle* sp. on red porgy [35] and the low clearance rates of albendazole (46.5% and 32.8%) against monogeneans in *Piaractus mesopotamicus* [36] demonstrate the challenge of achieving therapeutic drug concentrations across the monogenean tegument during rapid immersion. Although triclabendazole is highly effective against endoparasite flukes following oral administration [37], its application as a bath treatment for ectoparasitic monogeneans requires prolonged exposure or high concentrations [38], further emphasizing the pharmacokinetic barriers inherent to immersion therapy and the fundamental physiological differences between endoparasitic trematodes and ectoparasitic monogeneans.

Even among agents specifically targeted at monogeneans, conventional therapies face substantial limitations. Praziquantel (PZQ), widely regarded as a frontline treatment for *Neobenedenia* infections, achieves only partial and highly variable efficacy (19-86%) against *N. girellae* when administered orally [22,39]. In bath applications, Morales-Serna et al. (2018) [23] reported that 3 mg/L PZQ required 12 h to attain 87% mortality of *N. melleni* adults, while the commercial combination product Adecto® (20 mg/L) required 12-16 h to achieve 100% adulticidal activity. By comparison, compound D, a 5,6-dihydropyridine derivative bearing a tert-butyl carbamate and an ethyl ester, exhibited potent concentration-dependent antiparasitic efficacy, achieving 76.54% clearance at 0.5 mg/L and 95.23% at 0.7 mg/L following 90 min immersion. Crucially, this rapid action occurs within a tolerable host safety threshold: high *in vitro* cell viability (>80% in GF-1 cells) and clean histopathological profiles in groupers at effective bath concentrations (0.7 mg/L) confirm that compound D possesses intrinsic selectivity for parasitic targets over host tissues. Although acute mortality at higher concentrations (>1.0 mg/L) defines a narrow therapeutic window, these findings collectively demonstrate that compound D overcomes the trans-tegumental absorption barrier that limits traditional antiparasitic drugs, establishing it as a promising lead for immersion-based monogenean control.

The mechanism underlying this rapid contact toxicity appears to involve direct tegumental damage and potential cytoskeletal interference. Scanning electron microscopy revealed extensive surface folding, fissures, and structural disruption across the body surface of *N. melleni* following 0.7 mg/L compound D exposure, indicative of severe tegumental compromise. Such ultrastructural alterations are consistent with a membrane-active mode of action; however, the molecular basis of this damage warranted further investigation through target identification. The ten proteins selected for docking were not chosen arbitrarily. Each has been implicated as a drug target, vaccine candidate, or essential survival protein in parasitic helminths or protozoa: β-tubulin is the established target of benzimidazole anthelmintics in nematodes, cestodes and trematodes [40,41]; cathepsin B and cathepsin L-like cysteine proteases have been validated as key invasion and nutrient-acquisition enzymes in *Schistosoma* spp. and *Fasciola* spp. [42–44]; glutathione transferase and heat shock proteins 70/90 are associated with drug resistance and stress responses in multiple parasites [45,46]; and calcium-transporting ATPases serve as targets for antiparasitic compounds in apicomplexan parasites [47]. Because proteomic studies have detected homologues of these proteins in *Neobenedenia* sp. [29], they represent rational candidates for evaluating the potential mode of action of compound D in a monogenean context. Among these validated targets, β-tubulin showed by far the strongest predicted interaction with compound D (−6.53 kcal/mol, predicted Ki = 16.39 μM, three hydrogen bonds with MET255, ARG297, and LEU299), whereas cathepsin B, annexin B1 and glutathione transferase showed weaker binding (−3.12 to −3.67 kcal/mol), and the remaining proteins showed little predicted affinity. This hierarchy is informative: compound D appears to discriminate among established parasite targets, preferring the cytoskeletal protein over proteases, chaperones and ion transporters. The preferential docking to β-tubulin is also consistent with the tegumental disruption observed by SEM, because microtubule destabilisation is known to produce surface collapse and detachment in benzimidazole-treated helminths [48]. Whether compound D occupies the same binding pocket as benzimidazoles or an allosteric site on β-tubulin remains to be determined experimentally. Collectively, these mechanistic insights suggest that compound D may exert its antiparasitic effects through a dual action: rapid physical disruption of the tegumental barrier followed by interference with microtubule integrity, though further biochemical and functional studies are needed to confirm this hypothesis.

The present findings therefore identify compound D as a candidate for further development as an immersion treatment against *N. melleni*, rather than as a confirmed therapeutic agent at this stage. Several questions remain to be addressed, including the direct interaction between compound D and β-tubulin, its effects on microtubule organization and parasite viability, its toxicity following repeated or prolonged exposure, and its efficacy under larger-scale aquaculture conditions. Addressing these questions will be necessary to determine whether the promising activity observed here can be translated into a practical treatment strategy for *N. melleni* infection.

## 5. Conclusions

In conclusion, the present study identified the monogenean infecting cultured *Epinephelus fuscoguttatus*♀ × *E. lanceolatus*♂ as *Neobenedenia melleni* based on ITS1 sequence analysis and phylogenetic inference, providing a molecular basis for subsequent antiparasitic evaluation. Among six candidate compounds tested, compound D showed the highest antiparasitic efficacy against *N. melleni* and exhibited a clear concentration-dependent activity, reaching 95.23% efficacy at 0.7 mg/L after 90 min of immersion. At this concentration, compound D was well tolerated by groupers, with no apparent histopathological alterations in the examined tissues. Scanning electron microscopy further revealed severe surface folding, fissures, and structural disruption in treated parasites, indicating pronounced tegumental damage. Molecular docking against ten parasite-associated proteins identified β-tubulin as the protein with the most favorable predicted interaction with compound D, suggesting that β-tubulin may be a potential molecular target involved in its antiparasitic activity. Taken together, these findings identify compound D as a promising lead compound for the development of immersion-based control strategies against *N. melleni*. However, direct binding assays and functional studies of β-tubulin and microtubule organization, together with repeated-exposure toxicity and larger scale efficacy evaluations, are required to further validate its mode of action and practical potential in aquaculture.

## Author Contributions

Longkun Gao and Wei Luo: Writing – original draft, Visualization, Methodology, Investigation; Guangshuo Wang and Yanru Guo: Methodology, Formal analysis; Ying Yan, Guanhai Li and Juntao Xu: Methodology, Visualization; Qin Yu and Mingzhu Liu: Formal analysis; Erlong Wang: Formal analysis, Conceptualization; Pengfei Li: Resources, Conceptualization; Tianqiang Liu: Project administration, Funding acquisition, Writing – review & editing. All authors have read and agreed to the published version of the manuscript.

## Institutional Review Board Statement

All animal experiments were approved by the Northwest A&F University Animal Care and Use Committee, and we have complied with all relevant ethical regulations (DK20250512001).

## Data Availability Statement

Data are contained within the paper.

